# Non-responder phenotype reveals microbiome-wide antibiotic resistance in the murine gut

**DOI:** 10.1101/566190

**Authors:** Christian Diener, Anna C. H. Hoge, Sean M. Kearney, Susan E. Erdman, Sean M. Gibbons

## Abstract

Broad spectrum antibiotics can cause both transient and lasting damage to the ecology of the gut microbiome. Loss of gut bacterial diversity has been linked to immune dysregulation and disease susceptibility. Antibiotic-resistant populations of cells are known to arise spontaneously in single-strain systems. Furthermore, prior work on subtherapeutic antibiotic treatment in humans and therapeutic treatments in non-human animals have suggested that entire gut communities may exhibit spontaneous resistance phenotypes. In this study, we validate the existence of these community resistance phenotypes in the murine gut and explore how antibiotic duration or diet influence the frequency of this phenotype. We find that almost a third of mice exhibit whole-community resistance to a therapeutic concentration of the *β*-lactam antibiotic cefoperazone, independent of antibiotic treatment duration or xenobiotic dietary amendment. These non-responder (i.e. resistant) microbiota were protected from biomass depletion, transient ecological community collapse, and lasting diversity loss seen in the susceptible microbiota. There were no major differences between non-responder microbiota and untreated control microbiota at the community structure level. However, gene expression was vastly different between non-responder microbiota and controls during antibiotic treatment, with non-responder communities showing an upregulation of antimicrobial resistance genes and a down-regulation of central metabolism. Thus, non-responder phenotypes appear to combat antibiotic assault through a combination of efflux transporter upregulation and a reduced growth rate across the entire gut community. Future work should focus on what factors are responsible for tipping entire communities from susceptible to resistant phenotypes so that we might harness this phenomenon to protect our microbiota from exposure to therapeutic antibiotic treatment regimes.

## Introduction

Despite the clear public health utility of antibiotics, there is an undeniable cost to their widespread use in medicine and agriculture^1^. Antibiotic resistance in pathogens is on the rise and evidence is mounting that antibiotic treatments cause damage to our commensal microbiota^2–4^. The gut microbiome is an integral component of the human body, helping with nutrient absorption, pathogen resistance, and immune system education^2^. When the ecology of the gut is compromised by antibiotics, host health can suffer^5–7^.

Previous work in humans has shown that one round of antibiotic treatment can temporarily alter the taxonomic composition of the gut microbiome, increase the prevalence of antibiotic resistance genes, and lead to a permanent loss of species diversity^8–14^. The steady decline of gut bacterial diversity in developed nations over the last century, likely due in part to antibiotic use, has been implicated in the rise of chronic immune dysfunction^3,13,15^. Thus, finding ways to prevent or mitigate the ecological damage done by antibiotics is an important public health priority^13^. For example, strategies have been developed to introduce activated carbon into the lower gut during antibiotic exposure to protect colonic bacteria^16^ or to use autologous fecal transplants to replenish gut diversity following treatment^17^. In addition to these therapeutic strategies, the microbiome appears to exhibit natural antibiotic-resistance under certain conditions. Sup-therapeutic doses of antibiotics in animal models have been shown to substantially reduce gut microbiome diversity and biomass in some hosts but not others, indicating that these communities vary in their capacity for resistance^18–20^. In single-strain systems, sub-populations of antibiotic-resistant cells arise spontaneously due to stochastic apportionment of efflux transporters between daughter cells^21,22^ or due to the spontaneous amplification of antimicrobial resistance genes in mutant sub-populations^23^. Analogous symmetry-breaking processes^21,22,24^ may contribute to observed community-level antibiotic resistance in the microbiome^18,20^.

Heterogeneous responses of gut microbiota to therapeutic antibiotic treatments have been reported in the literature^10,18,25,26^. For example, while antibiotic exposure is a risk factor for *Clostridium difficile* carriage and infection in hospitals, not all antibiotic-treated patients exposed to *C. difficile* become infected^27,28^. To investigate this phenomenon in a controlled system, Schubert et al. (2015) looked into how the type and concentration of antibiotic treatment influenced *C. difficile* colonization of the murine gut^29^. The authors built a Random Forest (RF) regression model that could accurately predict *C. difficile* colonization levels from the composition of the gut microbiome. Cefoperazone, a broad-spectrum *β*-lactam antibiotic, had a large effect on the composition of the gut microbiome across most mice, lowered bacterial biomass by three orders of magnitude, and made mice susceptible to *C. difficile* colonization and infection^29^. Interestingly, several mice that received relatively high concentrations of cefoperazone were not colonized by the pathogen^29^. These resistant mice were also not predicted to be colonized by the RF model and thus appeared to maintain a gut microbiome similar in composition to the control mice. Based on these results, we hypothesized that whole-community antibiotic resistance to therapeutic levels of antibiotics might be a common phenomenon in the mammalian gut.

In this study, we explore the potential mechanisms underlying the cefoperazone non-responder phenotype (i.e. cefoperazone resistant microbiomes) and look at how the prevalence of this phenotype varies across treatment regimes. Although it was not a focus of their work, Schubert et al. (2015) showed that the frequency of non-responder phenotypes decreased with higher concentrations cefoperazone,^29^ which comports with prior work on sub-therapeutic antibiotic treatments in mice^18,19^. Other important factors that could influence the frequency of these non-responder phenotypes are the duration of antibiotic exposure^30^ and host diet^31,32^. We designed and carried out two independent mouse experiments to explore the reproducibility and frequency of non-responder phenotypes to a therapeutic dose of cefoperazone (100-175 mg/kg/day) across duration and dietary treatments. For the diet experiment, we included a 1% seaweed amendment to normal mouse chow, as used previously by our group^32^. We hypothesized that previous exposures to plant-derived secondary compounds might influence subsequent responses to antibiotics^33,34^, and raw seaweed is a rich source of these compounds^35,36^. In addition to measuring community composition and biomass, we sequenced community transcriptomes in non-responder and control microbiomes to characterize the gene expression profiles underlying community-wide resistance.

Overall, we found that 31% (i.e. 10 out of 32) of singly-housed mice exposed to therapeutic levels of cefoperazone were protected from antibiotic-induced collapse of the gut microbiome, independent of duration or dietary treatments. The community structure, species diversity, and biomass of non-responder microbiomes were similar to untreated controls and reproducible across both experiments. Despite little-to-no change in community composition, non-responder microbiota showed dramatic differences in community transcriptional profiles when compared to untreated mice (i.e. > 25% of all gene functions were differentially expressed). Gene functions involved in growth and motility were downregulated and antimicrobial resistance (efflux transporters, in particular) was upregulated in non-responder microbiomes. Together, these results indicate that entire bacterial communities can spontaneously protect themselves from collapse in the presence of a broad-spectrum antibiotic, likely through a combination of quiescence and antimicrobial resistance.

## Results and Discussion

### Antibiotic duration experiment

28 week old female C57BL/6J mice from the same birth cohort were cohoused (5-6 mice per cage) prior to beginning the experiment, and then separated into individual cages 1 week prior to antibiotic treatments. Singly-housed mice were exposed to 0.5 mg/mL^29^ cefoperazone in their drinking water for 0, 2, 4, 8, or 16 days (Fig. 1A). C57BL/6J mice drink an average of 6 mL of water each day^37^, so the dosage of cefoperazone was well within the therapeutic range (100-150 mg/kg/day). 16S amplicon sequencing showed the majority of cefoperazone-treated mice experienced dramatic restructuring of their microbiome composition in response to antibiotics (Fig. 1B). However, 6 of the 16 cefoperazone-treated mice in this duration experiment did not exhibit a drastic change in microbiome composition over the course of the experiment. Thus, the microbiota in these mice were resistant. The only duration treatment where all mice responded to antibiotic treatment was the 4-day duration. Overall, duration of exposure had no significant influence over the frequency of non-responder phenotypes (Fisher’s Exact Test p=0.44). The microbiome composition of non-responder mice was indistinguishable from untreated control mice at the phylum level (Fig. 1C), but there were detectable differences in Bacteroides, Akkermansia, and Lachnospiraceae at the amplicon sequence variant (ASV) level (see Methods and Fig. 1B). Susceptible mice showed a major turnover in community composition at the phylum level, with a near-complete loss of Bacteroides and Firmicutes and a dramatic enrichment of Proteobacteria and Cyanobacteria. Proteobacteria and Cyanobacteria reads were identified as being derived largely from mitochondria (likely from host) and chloroplasts (likely from plant-based diet), respectively. Initially, we had predicted that duration of exposure would be positively correlated with ASV extinction rates (i.e. ASVs present within a mouse initially, but not at the end of the experiment). Treatment duration had no significant effect on loss, gain (i.e. absent initially within a mouse, but present at the end of the experiment), or persistence (i.e. present within a mouse at the beginning and end of the experiment) of ASVs across the entire data set (ANOVA p > 0.1). There was a clear significant increase in species extinctions and a decrease in persistent ASVs in antibiotic susceptible mice compared to control mice (Fig. 1D). Non-responder mice, however, showed no significant differences from controls in ASVs gained, lost, or persistent (Fig. 1D). Thus, non-responder microbiota were protected from phylum-level collapse of gut bacterial community structure following antibiotics and from antibiotic-associated diversity loss.

**Figure 1.**
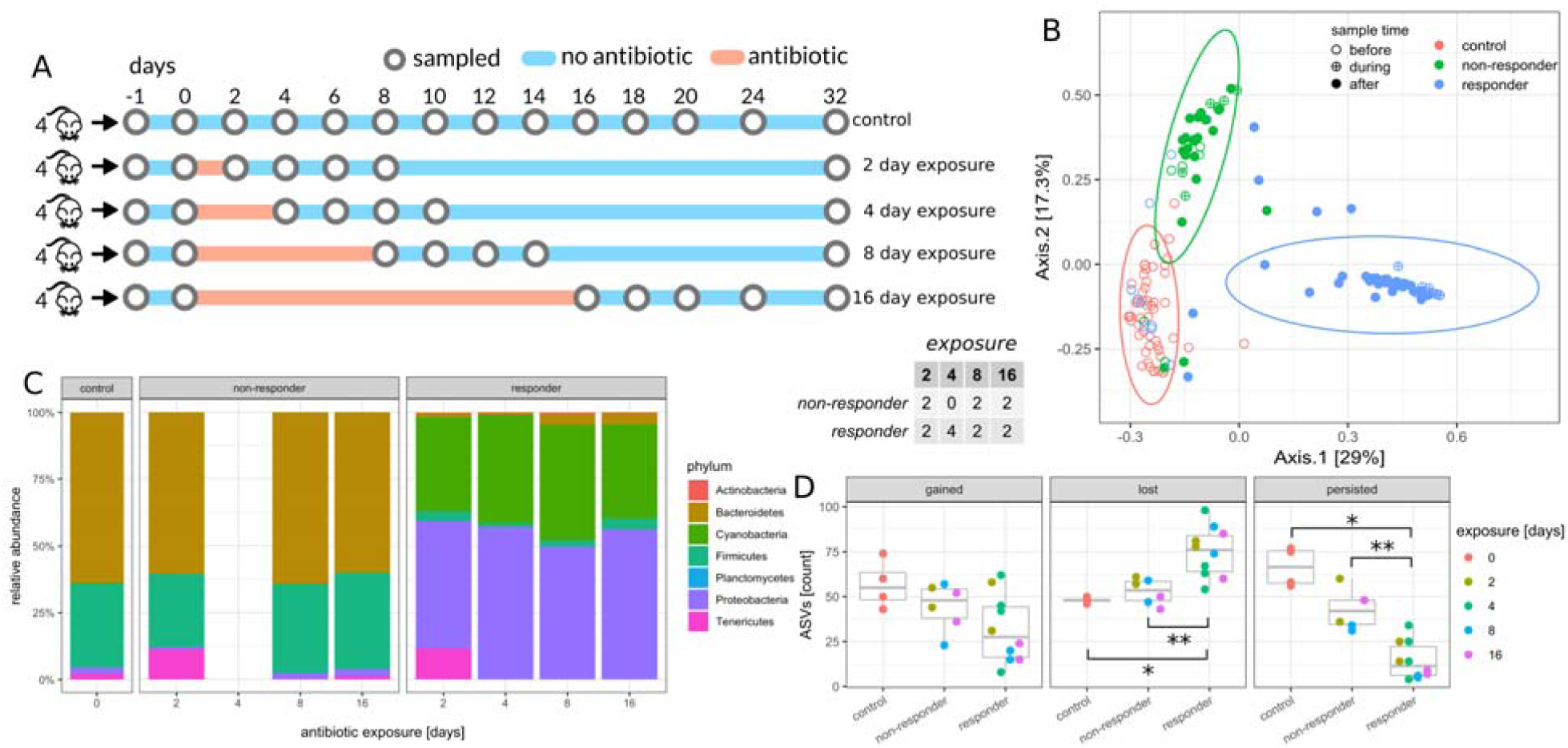
Effect of antibiotic exposure duration on non-responder phenotype. Table in the center denotes number of non-responder and responder mice in each treatment duration group. (A) Experimental design for the duration experiment. Circles denote sampled time points. (B) Principal coordinate analysis (PCoA) of samples during and after antibiotic exposure (n=143, day >= 0). Ellipses denote 95% confidence intervals from a Student t-distribution. Each point denotes a sample and annotated numbers denote days after antibiotics treatment. ASV abundances were rarefied to 10K reads for each sample and percentages in brackets denote the explained variance. (C) Relative abundance of phyla at the last day of antibiotics treatment. The control panel is an average over all untreated controls. Only phyla with a relative abundance of at least 0.1% are shown. (D) Dynamics of amplicon sequence variants (ASVs). Gained ASVs are variants that were not present before antibiotics treatment but are present after. Similarly, lost ASVs were present before treatment but not after, and persistent ASVs were present before and after.

### Seaweed diet experiment

14 seven-week-old female C57BL/6J mice from the same birth cohort were cohoused prior to beginning the experiment (5-6 mice per cage), and then separated into individual cages 1 week prior to dietary treatments. Half of the mice were given a 1% seaweed in normal chow diet and the other half received a normal chow diet for 20 days (Fig. 2A). All mice were put on a normal chow diet for six days prior to antibiotic treatment. On day 26, all mice continued on a normal chow diet and replicate mice from each dietary treatment group were given 0.5 mg/mL cefoperazone in their drinking water for a period of 6 days (Fig. 2A). On most sampling days, only the first three replicates from each treatment group were sampled for 16S sequencing.

**Figure 2.**
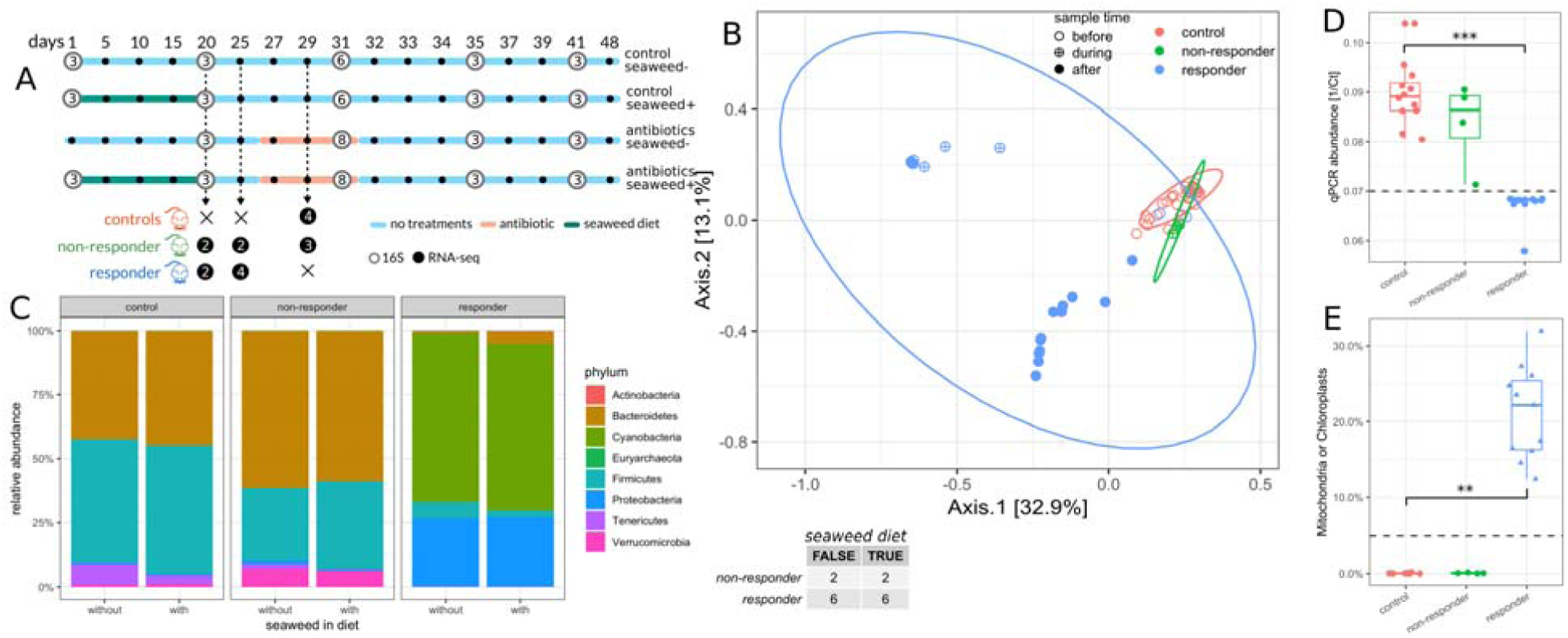
Effect of seaweed diet on non-responder phenotype. Table in the center denotes number of non-responder and responder mice in each diet group. (A) Design of the diet experiment. White circles denote 16S samples and are filled with the number of biological replicates for each sampling point. Black circles denote RNA-seq samples. (B) PCoA of 16S samples after diet (n=60, day >= 20). Symbol fill denotes sampling time relative to antibiotic treatment. Ellipses denote 95% confidence interval from a Student t-distribution. ASV abundances were rarefied to 5K reads for each sample and percentages in brackets show explained variance. (C) Relative phyla abundances across diet and response groups. Only phyla with a relative abundance larger than 0.1% are shown. (D) qPCR biomass estimates (1/Ct) for samples across response groups. (E) Percentage of mitochondria and chloroplast sequences in 16S amplicon data across response groups. Triangles indicate samples below dashed line in panel D, considered to be low-biomass, while circles indicate high biomass samples

However, on the final day of antibiotic treatment, the full set of replicate mice were sampled (16 antibiotic-treated mice and 12 control mice). Seaweed treatment had a very minor impact on the composition and diversity of the microbiome (Fig. S1), similar to what we had observed previously^32^. We found the same non-responder and responder phenotypes as in the duration experiment, with 4 of the 16 mice exhibiting the non-responder phenotype (Fig. 2B-C). The seaweed diet had no effect on the frequency of the non-responder phenotype (Fisher’s Exact Test p = 1.0). We measured total 16S copy numbers for each sample (i.e. a proxy for bacterial biomass) and found that antibiotic susceptible mice showed a dramatic drop in fecal bacterial biomass following cefoperazone treatment, while non-responder microbiomes did not differ significantly from controls in biomass levels (Fig. 2D). Similar to the first study, we saw an enrichment in mitochondrial and chloroplast sequences in the susceptible mice, which also corresponded to the drop in bacterial biomass (Fig. 2D-E). Thus, it appears that the absence of appreciable bacterial biomass in a mouse stool results in an enrichment for host and dietary contaminants in 16S amplicon sequencing data.

### Temporal dynamics following antibiotic treatment

Despite greater loss of species in susceptible mice (Fig. 1D), overall alpha diversity tended to recover over time in these mice after cessation of antibiotic treatment. Non-responder and control mice maintained relatively stable alpha diversities over time, although antibiotic-treated, non-responder microbiota showed slightly lower alpha diversities than controls (Fig. 3A). Despite the resilience of Shannon diversity in the susceptible mice over time, only a small number of these mice showed recovery in Bacteroidetes ASVs (Fig. 3B). In control and non-responder mice, Bacteroidetes was the dominant phylum over the entire time series. However, Firmicutes became the dominant phylum in responder mice following antibiotics and in many mice there appeared to be a permanent loss of the Bacteroidetes phylum following recovery (Fig. 3B). Seaweed dietary treatment appeared to contribute to a loss in resilience, with none of the seaweed-fed susceptible mice showing recovery of the Bacteroides phylum (Fig. S1).

**Figure 3.**
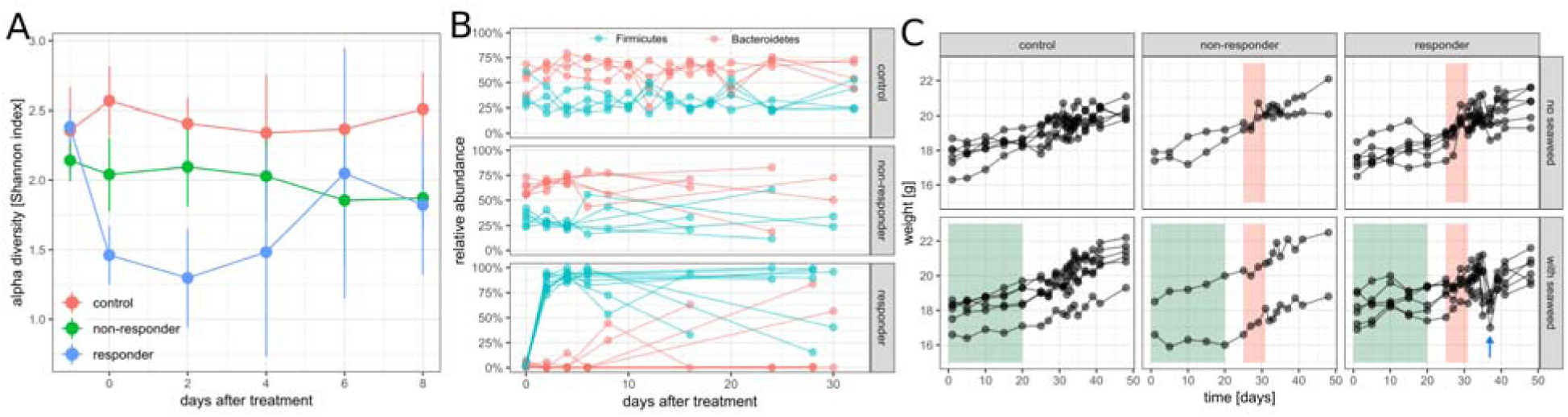
Temporal dynamics in non-responder and responder mice following antibiotic and diet treatments. (A) Alpha diversity (Shannon index) dynamics after antibiotics treatment in the duration experiment. Each point denotes mean of all samples regardless of duration and error bars denote standard deviation. (B) Dynamics of Bacteroidetes and Firmicutes phyla in the antibiotic duration experiment. (C) Mouse weights in the diet experiment. Green areas denotes seaweed diet treatment windows and red area denotes antibiotics treatment windows. The blue arrow indicates transient weight loss in seaweed-fed mice a few days following the end of antibiotic treatment.

Post-antibiotics, the susceptible microbiomes were dominated by Firmicutes (Figure 3B). In the diet experiment, in which mice were sampled on the last day of antibiotics and then 4 days post-antibiotics, most Firmicutes ASVs belonged to the order Clostridiales. In the duration experiment, however, mice were sampled 2 days post-antibiotics. At the 2-day timepoint, some Firmicutes populations were dominated by ASVs from the order Lactobacillales, while the rest showed the familiar Clostridiales-dominated signature. By 4-days post-antibiotics, however, Clostridiales had reached greater than 82% relative abundance in all mice, and the Lactobacillales population had uniformly dwindled to less than 3.4% relative abundance. Thus, Lactobacillales ASVs appear to be involved in rapid recovery following cessation of antibiotic treatment, but are quickly displaced by Clostridiales ASVs.

Despite the fact that we saw no major differences in the gut microbiomes between control diet (i.e. normal chow) mice and 1% seaweed-fed mice (Figs. 2C and S1), we did observe a difference in mouse weight loss (Fig. 3C). All antibiotic susceptible mice that were fed seaweed showed a large, transient weight-drop a few days following the cessation of antibiotics (Fig. 3C). Antibiotic-treated mice that did not receive seaweed also showed a mild drop in weight, but the effect was weaker (Fig. 3C). This weight loss phenotype was not observed in control mice that were not treated with cefoperazone and was also not observed in antibiotic non-responder mice from both diet treatment groups (Fig. 3C). We do not have an explanation for this synergistic effect between seaweed diet and cefoperazone treatment on transient weight loss in mice, but we believe this to be an interesting research avenue to explore further.

### Non-responder microbiomes exhibit an antimicrobial resistance transcriptional program

To evaluate whether the occurrence of the non-responder phenotype might be associated with changes in gene transcription in the gut, we performed RNA sequencing on samples from 10 mice before antibiotic treatment (days 20 and 25) and 7 mice during antibiotic treatment (day 29). Because there was close to no biomass in responder fecal samples after antibiotics treatment, we compared non-responders to untreated controls on day 29. After RNA extraction, ribosomal depletion, and sequencing to a mean of 20 million reads per sample, we identified around 800,000 unique transcripts by *de novo* assembly (see Material and Methods) ranging from 111 to >26,000 bp lengths (longer contigs were polycistronic; see Fig. S2 for length and coverage distributions). Non-summarized transcript abundances were sufficient to distinguish non-responder communities from controls after antibiotic treatment (Fig. 4A). However, that may be due to the high specificity of transcripts for each sample since we found that, on average, each transcript only appeared in 3 of the 17 total samples (Fig. S3A). Thus, we also also assigned functional annotations to transcripts by aligning them to the M5NR database^38^. We were able to assign 61% of the original transcripts to functions in the SEED subsystem database^39^. This allowed us to collapse transcript counts for each sample into SEED functions, which yielded a total of 53,877 unique functions. The majority of SEED functions were detected in all 17 RNA-seq samples (see Fig. S3B). Control and non-responder communities could be easily distinguished by functional counts (see Fig. 4B). In particular, about 45% of the variance in functional expression could be explained by non-responder vs. control status (Euclidean PERMANOVA p = 0.036) compared to only 6.7% of explained variance in 16S beta diversity (Bray-Curtis PERMANOVA p = 0.027). Thus, transcriptional differences capture the non-responder phenotype much better than changes in community composition.

**Figure 4.**
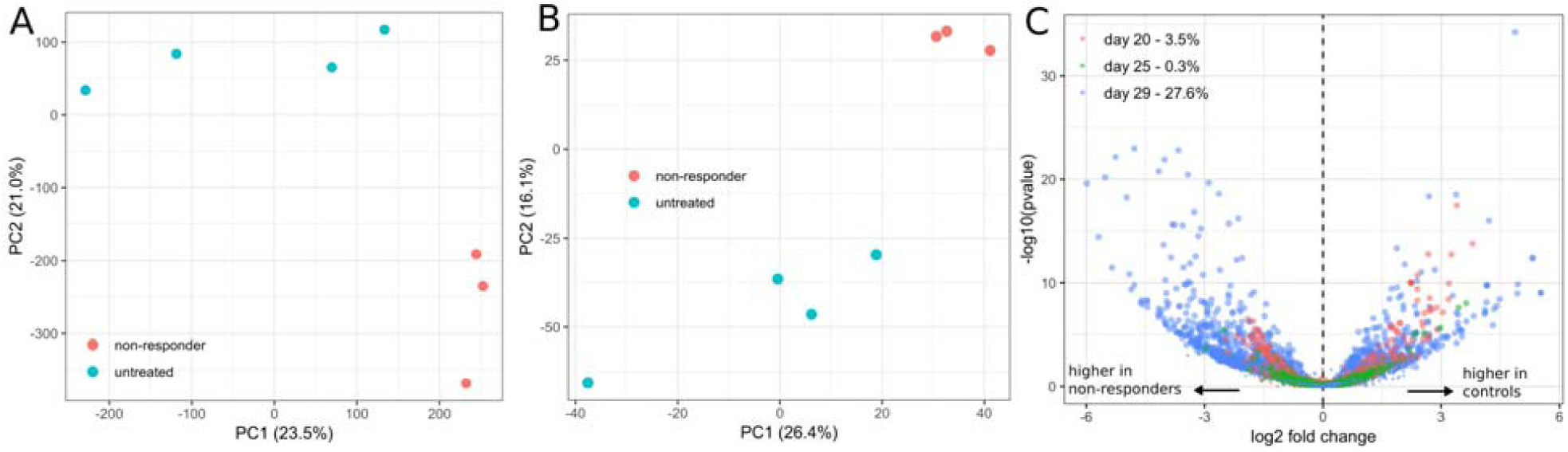
Global transcriptional response to antibiotics in non-responder phenotypes. (A) Principal component analysis (PCA) of post-treatment samples based on abundances of *de novo* assembled transcripts. Percentages in brackets denotes explained variance. (B) PCA of post-treatment samples based on abundances of functional groups. (C) Volcano plot of untreated vs. non-responder differential abundance tests (functional groups). Percentages for each day denote positive tests rate (number of significant tests / total tests) and colors denote day the samples were taken (20 and 25 were before antibiotic treatment and day 29 was after). Tests with FDR q-values < 0.05 are shown as larger dots, whereas non-significant results are shown as small dots.

After filtering out low abundance functions, differential expression testing between controls and non-responder communities was performed for each of the three time points sampled (see Materials and Methods). We observed that less than 5% of the observed functions were differentially expressed at an FDR q ≤ 0.05 before antibiotic exposure (days 20 and 25), which fits our null-expectation. However, following antibiotic exposure (day 29), 27% of all functions were differentially expressed between untreated controls and non-responder communities (see Fig. 4C). This indicated a global transcriptional shift in non-responder microbiomes, mostly characterized by a up-regulation of several functional groups in the non-responder mice (blue dots on left side of Fig. 4C).

The transcriptional program was most prominently characterized by an upregulation of efflux transporters and other antibiotic resistance defense mechanisms, and a down-regulation of motility and respiratory functions. For instance, the SEED sub-pathway “Transporters in Models” was the most prominent subpathway in the differentially expressed functions, containing 82 significant hits (FDR q ≤ 0.05). Most of the significantly upregulated functions in the “Virulence, Disease and Defense” superpathway were also related to efflux pumps and their regulation (Fig. 5A). We also found large differences in of respiratory pathways, albeit with a mixed pattern of up- and down-regulation (Fig. 5B). The most striking respiratory difference we observed was the down-regulation of three acetyl-CoA synthases which were some of the most highly expressed functions in the untreated mice (Fig. 5E). These pathways were down-regulated by one to two orders of magnitude in the non-responder mice, which suggests a down-regulation of central metabolism. Additionally, we observed a consistent down-regulation of flagellar motor proteins in the non-responder mice (Fig. 5C). All differentially expressed functions in the “Membrane Transport” superpathway were strongly upregulated in the non-responder mice, including components of TonB, which is known to be necessary for efflux transporter function^40^ (Fig. 5D). Together, these data are consistent with previous reports that upregulation of efflux transporters is accompanied by a concomitant reduction in growth rate^21,41^. Finally, we observed the upregulation of the entire vancomycin resistance locus, including the three efflux pumps Vex1-3 and the two-component system VncR and VncS (Fig. 5E). The induction of vancomycin cross-resistance by *β*-lactams has been described before^42,43^ and might indicate that these loci confer general efflux-based resistance to a range of antibiotics.

**Figure 5.**
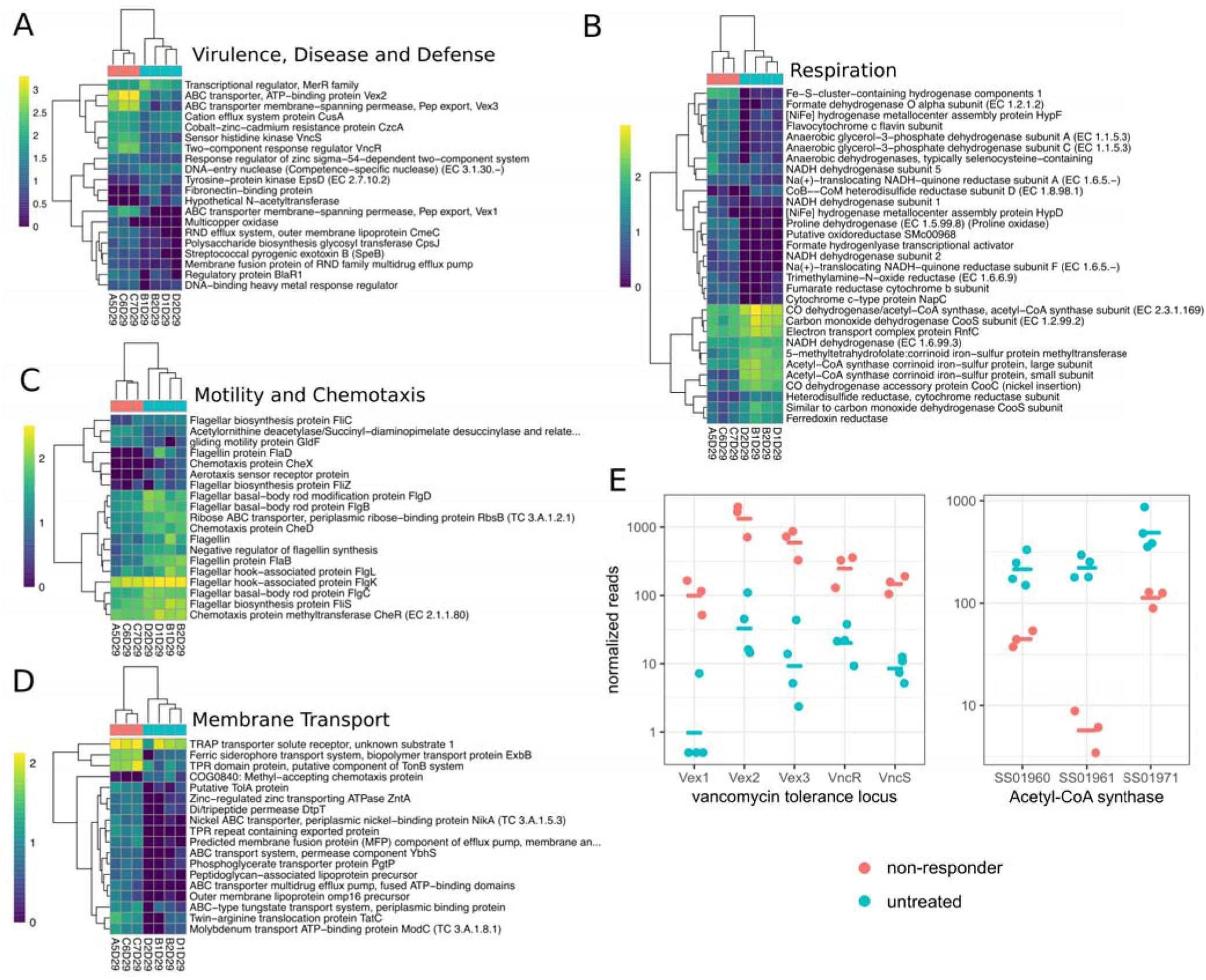
Differentially abundant pathways in non-responder phenotypes. (A-D) Heatmaps showing differentially abundant (FDR < 0.05) functional groups grouped by SEED superpathway. Heatmap color scale shows normalized reads on a log10 scale with a pseudo count of 1. (E) Normalized expression of genes on the vancomycin tolerance locus and three Acetyl-CoA synthase genes between non-responders and controls.

## Conclusion

We found that nearly one third of mice exposed to therapeutic levels of the *β*-lactam antibiotic cefoperazone were protected from gut microbiome community turnover, biomass collapse, and diversity loss. The frequency of this non-responder phenotype does not depend on duration of antibiotic exposure or on seaweed being added to the diet, but does appear to increase as the concentration of cefoperazone in drinking water declines, as shown previously^29^. Despite very minor changes in community composition and diversity between untreated and non-responder mice, we observe a striking difference in microbiome gene expression between these groups of mice. Non-responder microbiomes show down-regulation of central metabolism and motility and upregulation of antimicrobial resistance. This combination of increased resistance and quiescence appears to protect gut communities from the extensive ecological damage observed in antibiotic-susceptible microbiomes. While prior work has shown how isogenic sub-populations of cells and two-species communities can exhibit heterogeneous responses to antibiotics^20–23^, the exact mechanisms underlying transitions into whole-community resistance phenotypes in the mammalian gut are not yet clear and will require further study. Future work should focus on what factors tip microbiomes between non-responder and responder phenotypes, potential hysteresis of these phenotypes, and whether or not this transition point can be manipulated to protect commensal microbiota from antibiotic assault.

## Materials and Methods

### Animal care

5- to 6-week-old female C57BL/6J mice ordered from Jackson Laboratories (Bar Harbor, ME) were housed were housed and handled in Association for Assessment and Accreditation of Laboratory Animal Care (AAALAC)-accredited facilities using techniques and diets specifically approved by Massachusetts Institute of Technology’s Committee on Animal Care (CAC) (MIT CAC protocol no. 0912-090-15 and 0909-090-18). The MIT CAC (Institutional Animal Care and Use Committee [IACUC]) specifically approved the studies as well as the single-housing and handling of these animals. Mice were euthanized using carbon dioxide at the end of the experiment.

### Antibiotic duration experiment

For this 34-day experiment, 20 mice were assigned randomly and evenly to 5 treatment groups: control, 2 days of antibiotic exposure, 4 days of antibiotic exposure, 8 days of antibiotic exposure, and 16 days of antibiotic exposure. The *β*-lactam antibiotic cefoperazone was administered through drinking water at a concentration of 0.5 mg/mL, as in prior work^29^. Fecal samples were collected on the 2 days preceding antibiotic exposure, the last day of antibiotic exposure, and select timepoints following antibiotic exposure. Mice were weighed each sampling day. Fresh fecal samples were obtained within an hour of one another each day from all animals. Fecal samples were collected into 2 mL freezer tubes with 100 uL of anaerobic 40% glycerol containing 0.1% cysteine and transferred immediately to dry ice before being stored at −80° C prior to nucleic acid extraction.

### Seaweed diet and antibiotic experiment

A new cohort of 28 mice were split randomly into two diet treatment groups and were fed with either a custom chow diet (Bio-Serv, Flemington NJ) containing 1% raw seaweed nori (Izumi Brand) or a standard control diet (product no. F3156; AN-93G; Bio-Serv, Flemington NJ). Prior to the experiment, animals were co-housed for 10 days and then singly housed for 7 days prior to separation into the seaweed treatment and control groups. After 20 days of dietary treatment, all mice resumed the standard diet. From day 26 to day 31, 8 mice from each diet group were administered 0.5 mg/mL cefoperazone in their drinking water as in the duration experiment. Mice were weighed daily. Fresh fecal samples were obtained within an hour of one another each day from all animals in all groups. Fecal samples were collected into anaerobic 40% glycerol containing 0.1% cysteine and transferred immediately to dry ice before being stored at – 80° C prior to nucleic acid extraction.

### 16S amplicon sequencing

#### DNA extractions

DNA from fecal samples and bacterial cultures was extracted using the MoBio High Throughput (HTP) PowerSoil Isolation Kit (MoBio Laboratories; now QIAGEN) with minor modifications. Briefly, samples were homogenized with bead beating and then 50 μL Proteinase K (QIAGEN) added, and samples were incubated in a 65°C water bath for 10 min. Samples were then incubated at 95°C for 10 min to deactivate the protease.

#### Amplicon sequencing library preparation and biomass quantification

Libraries for paired-end Illumina sequencing were constructed using a two-step 16S rRNA PCR amplicon approach as described previously with minor modifications^44^. The first-step primers (PE16S_V4_U515_F, 5’-ACACG ACGCT CTTCC GATCT YRYRG TGCCA GCMGC CGCGG TAA-3’; PE16S_V4_E786_R, 5’-CGGCA TTCCT GCTGA ACCGC TCTTC CGATC TGGAC TACHV GGGTW TCTAA T-3’) contain primers U515F and E786R targeting the V4 region of the 16S rRNA gene, as described previously^44^. Additionally, a complexity region in the forward primer (5’-YRYR-3’) was added to help the image-processing software used to detect distinct clusters during Illumina next-generation sequencing. A second-step priming site is also present in both the forward (5’-ACACG ACGCT CTTCC GATCT-3’) and reverse (5’-CGGCA TTCCT GCTGA ACCGC TCTTC CGATC T-3’) first-step primers. The second-step primers incorporate the Illumina adaptor sequences and a 9-bp barcode for library recognition (PE-III-PCR-F, 5’-AATGA TACGG CGACC ACCGA GATCT ACACT CTTTC CCTAC ACGAC GCTCT TCCGA TCT-3’; PE-III-PCR-001-096, 5’-CAAGC AGAAG ACGGC ATACG AGATN NNNNN NNNCG GTCTC GGCAT TCCTG CTGAA CCGCT CTTCC GATCT-3’, where N indicates the presence of a unique barcode).

Real-time qPCR before the first-step PCR was done to ensure uniform amplification, avoid overcycling templates, and to provide a basic estimate of bacterial biomass for each sample (i.e. total copies of the 16S gene per volume of DNA extraction from a single mouse fecal pellet). Both real-time and first-step PCRs were done similarly to the manufacturer’s protocol for Phusion polymerase (New England BioLabs, Ipswich, MA). For qPCR, reactions were assembled into 20 μL reaction volumes containing the following: DNA-free H_2_O, 8.9 μL; high fidelity (HF) buffer, 4 μL; dinucleotide triphosphates (dNTPs), 0.4 μL; PE16S_V4_U515_F (3 μM), 2 μL; PE16S_V4_E786_R (3 μM), 2 μL; BSA (20 mg/mL), 0.5 μL; EvaGreen (20×), 1 μL; Phusion, 0.2 μL; and template DNA, 1 μL. Reactions were cycled for 40 cycles with the following conditions: 98°C for 2 min (initial denaturation); 40 cycles of 98°C for 30 s (denaturation); 52°C for 30 s (annealing); and 72°C for 30 s (extension). Samples were diluted based on qPCR amplification to the level of the most dilute sample and amplified to the maximum number of cycles needed for PCR amplification of the most dilute sample (18 cycles, maximally, with no more than 8 cycles of second-step PCR). For first-step PCR, reactions were scaled (EvaGreen dye excluded; water increased) and divided into three 25-μL replicate reactions during both first- and second-step cycling reactions and cleaned after the first and second step using Agencourt AMPure XP-PCR purification (Beckman Coulter, Brea, CA) according to manufacturer instructions. Second-step PCR contained the following: DNA-free H_2_O, 10.65 μL; HF buffer, 5 μL; dNTPs, 0.5 μL; PE-III-PCR-F (3 μM), 3.3 μL; PE-III-PCR-XXX (3 μM), 3.3 μL; Phusion, 0.25 μL; and first-step PCR DNA, 2 μL. Reactions were cycled for 10 cycles with the following conditions: 98°C for 30 s (initial denaturation); 10 cycles of 98°C for 30 s (denaturation); 83°C for 30 s (annealing); and 72°C for 30 s (extension). Following second-step clean-up, product quality was verified by DNA gel electrophoresis and sample DNA concentrations determined using Quant-iT PicoGreen dsDNA Assay Kit (Thermo Fisher Scientific). The libraries were multiplexed together and sequenced using the paired-end with 250-bp paired-end reads approach on the MiSeq Illumina sequencing machine at the BioMicro Center (Massachusetts Institute of Technology, Cambridge, MA).

#### 16S amplicon sequencing data analysis

Amplicon sequencing data was processed using DADA2^45^ and a custom 16S analysis pipeline available at https://github.com/gibbons-lab/mbtools. After performing general quality assessment, raw reads were filtered using the “filterAndTrim” method from DADA2 using a left trim of 10bp to avoid low complexity 5’ sequences and a maximum of 2 expected errors per read under Illumina model. Length truncation was performed based on the quality profiles and ensuring that sufficient overlap for merging remained. Reads in the duration experiment were truncated at 240 and 150 bps for forward and reverse reads respectively, and reads in the diet experiment were truncated at 240 and 170 bps. More than 88% of the reads in the duration experiment and 93% of the reads in the diet experiment passed quality filtering and were passed on to downstream processing with DADA2. Error rates were learned on a sample of 250 million bases and most of the inferred sequence variants could be merged across forward and reverse reads (>95% of preprocessed reads remaining). Less than 7% of all reads from both experiments were classified as chimeric and removed as well. Taxonomy was assigned to the sequence variants using the DADA2 Naive Bayes classifier with a bootstrap agreement of >50% and using the SILVA ribosomal database^46^ version 132. Species were assigned by exact alignment where possible. PERMANOVA was performed using the Bray-Curtis distance on rarefied read counts with the “adonis” function from the “vegan” package (https://CRAN.R-project.org/package=vegan). Amplicon sequence variants contributing to the separation of variances were identified from the coefficients of individuals regressions against the target variable (returned by the “adonis” function as well).

All workflows (as R notebooks), installation instructions and additional metadata are provided at https://github.com/gibbons-lab/mouse_antibiotics and allow reproduction of all results and figures from the manuscript starting from the raw data. More complex functionality is provided in a dedicated R package (“mbtools”) which is provided along with documentation at https://github.com/gibbons-lab/mbtools. Raw sequencing data can be found on the Sequence Read Archive (SRA) (https://www.ncbi.nlm.nih.gov/sra) under accession numbers SRRXXXXXX, SRRXXXXX, and SRXXXXXX (Bioproject ID BXXXXXXXX) [RAW DATA WILL BE MADE PUBLIC PRIOR TO PUBLICATION].

### RNA extraction, RNA sequencing, and RNAseq data analysis

#### RNA extraction

RNA was extracted from a total of 17 samples from Experiment 2 with the AllPrep PowerFecal DNA/RNA Kit (Qiagen USA, Cat. No. 80244). The 17 samples included 7 target samples taken during antibiotic treatment (4 untreated and 3 non-responder) and 10 negative controls prior to antibiotic treatment (day 20 and 25, 6 susceptible and 4 non-responder). RNA integrity numbers (RIN) were obtained using a 2100 BioAnalyzer (Agilent USA) with the Eukaryote Total RNA Nano Series II chip (Agilent USA). The majority of samples showed RINs above 5 and samples with lower RIN (5 of the control samples) were included in sequencing while controlling for the effect of low integrity in downstream analyses by explicitly including RIN as a confounder, as described previously^47^.

#### Library preparation and RNA sequencing

Ribosomal RNA was depleted from the 17 RNA-seq samples using the Ribo-Zero Gold rRNA Removal Kit (Illumina USA, Cat. No. MRZE724) and final concentrations were measured using the Qubit RNA HS Assay Kit (ThermoFisher Scientific USA, Cat. No. Q32852). Library preparation was performed using the TruSeq Stranded mRNA LT Sample Prep Kit (Illumina USA, Cat. No. RS-122-2101) and all samples were sequenced in single end mode in one run on an Illumina NextSeq (NS500720) for 85 cycles, which yielded a total of 464 million reads.

#### Data analysis

Raw sequencing reads were quality filtered using the “filterAndTrim” function from DADA2 with a left trim of 5bp and a maximum expected error (maxEE) of 1. More than 95% of the raw reads passed those filters and were used for all downstream analyses. No length truncation was performed due to the short length and high 3’ quality scores of the reads.

Transcripts were assembled *de novo* from the filtered reads with RNA Spades (version 3.12.0) across the full set of reads (http://cab.spbu.ru/software/rnaspades/)^48^ using the default parameters. Transcript abundances for each sample were quantified by aligning the filtered reads to the assembled transcripts with Bowtie2 version 2.3.4.3^49^. Mapping of unique reads to several transcripts was resolved by allowing up to 60 alternative alignments per read and counting the transcript abundances with an transcript length-aware Expectation-Maximization algorithm as used by Kallisto^50^.

Functional annotations for the *de novo* assembled transcripts were obtained by first aligning the transcripts to the M5NR database^38^ using DIAMOND version 0.9.21^51^. Functional annotations were then obtained by using the existing mapping between M5NR and the SEED subsystems database^39^ as downloaded from the MG-RAST FTP (ftp://ftp.metagenomics.anl.gov/data/misc/JGI/). Finally, abundances for functional groups were calculated by summing the reads for each unique SEED subsystem ID in each sample.

Normalization, differential abundance testing and false discovery rate (FDR) adjustment for assembled transcripts or functional groups were performed using DESeq2 version 1.18.1^52^. To avoid a bimodal p-value histogram, this was preceded by a prefiltering step removing features with an average abundances <10 reads or not appearing in at least two of the samples.

## Acknowledgements

CD, ACHH, and SMG were supported by the Washington Research Foundation Distinguished Investigator Award and by startup funds from the Institute for Systems Biology. SK and SMG were supported by the Center for Microbial Informatics and Therapeutics at MIT. Thanks to Eric Alm, Sui Huang, Nathan Price, Nitin Baliga, Naeha Subramanian, and members of the Gibbons Lab for helpful feedback on this work.

## Supplemental Material

**Figure S1.**
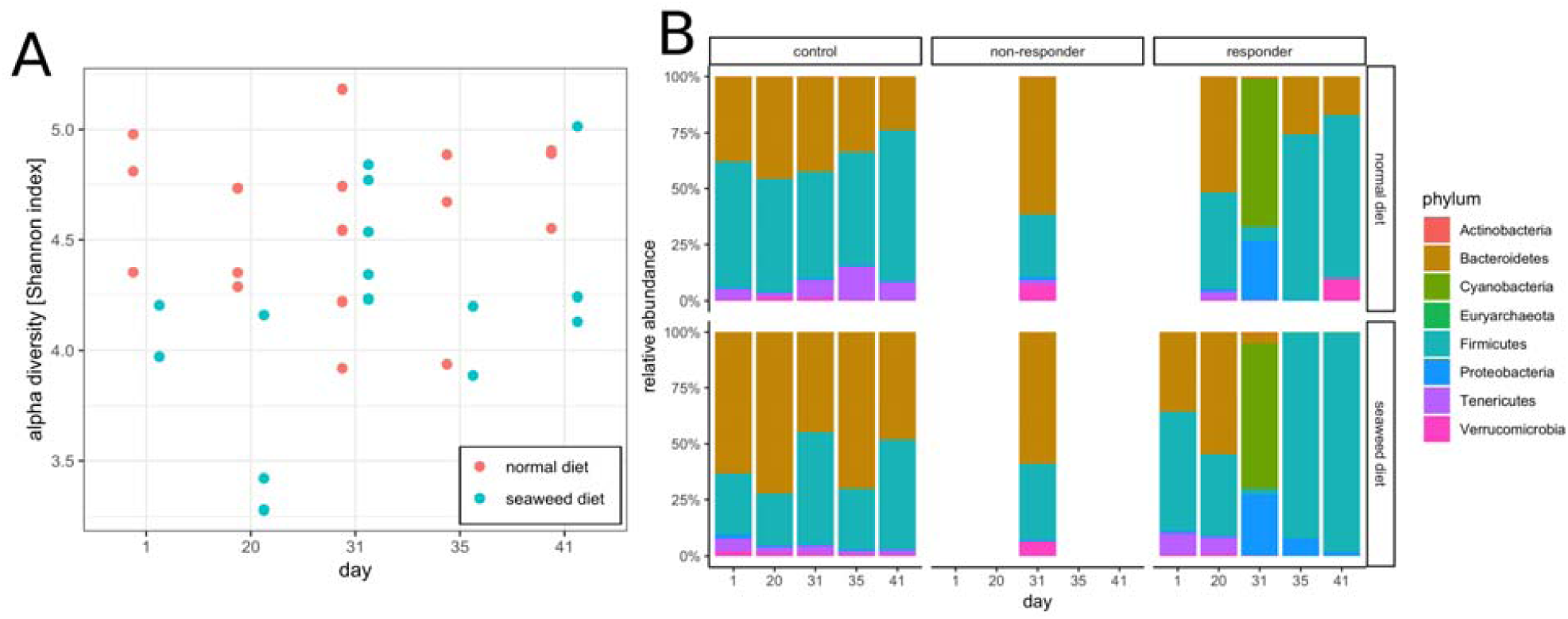
Alpha and beta-diversity trends for the diet study. (A) There were no significant differences in alpha diversity (Shannon index) over time between control mice (no antibiotics) fed a normal diet and mice fed a 1% seaweed diet for 20 days (Wilcoxon rank sum p > 0.1, for each day). All rarefied to 10000 reads each. (B) Community composition of control mice was slightly influenced by seaweed diet (7.7% explained variance, PERMANOVA p = 0.01) and this was mostly due to higher Bacteroides/Firmicutes ratio in mice fed the seaweed diet. Susceptible mice showed little differences before treatment but Bacteroidetes were completely lost in the seaweed treated susceptible mice (8% explained variance, PERMANOVA p = 0.01). Resistant mice did not show differences based on diet.

**Figure S2.**
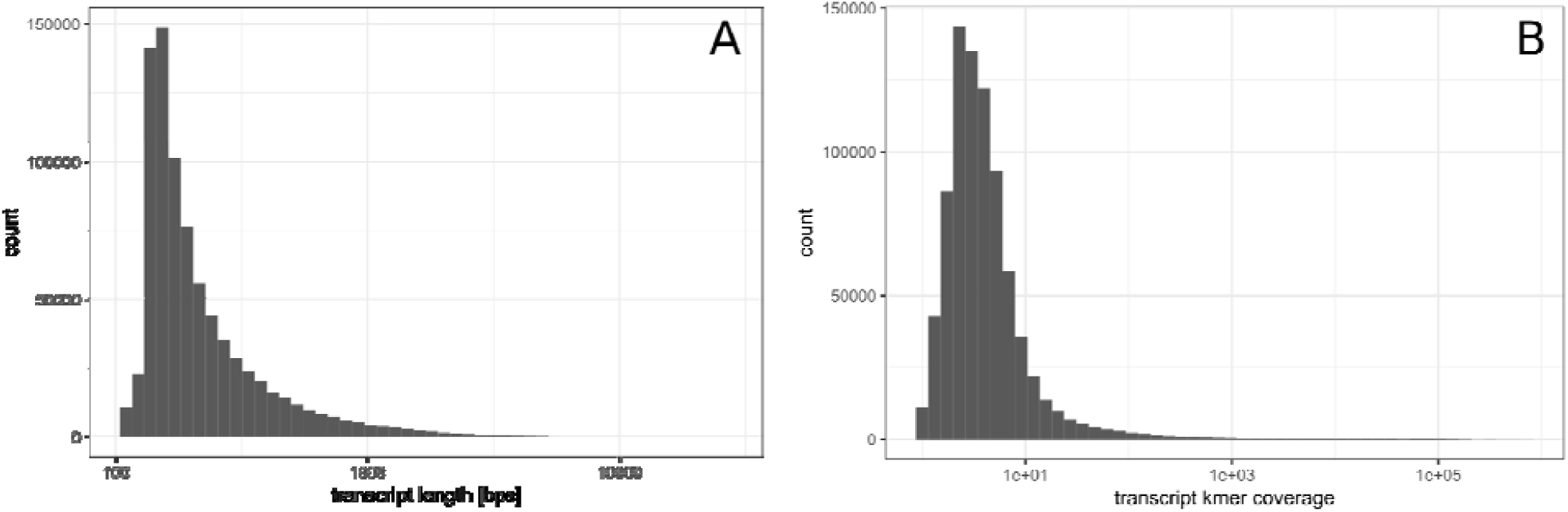
*De novo* assembly summaries for transcripts. (A) Length distribution of assembled transcripts. (B) Approximate coverage for assembled transcripts as estimated from k-mer coverage. Real coverage will always be larger than k-mer coverage.

**Figure S3.**
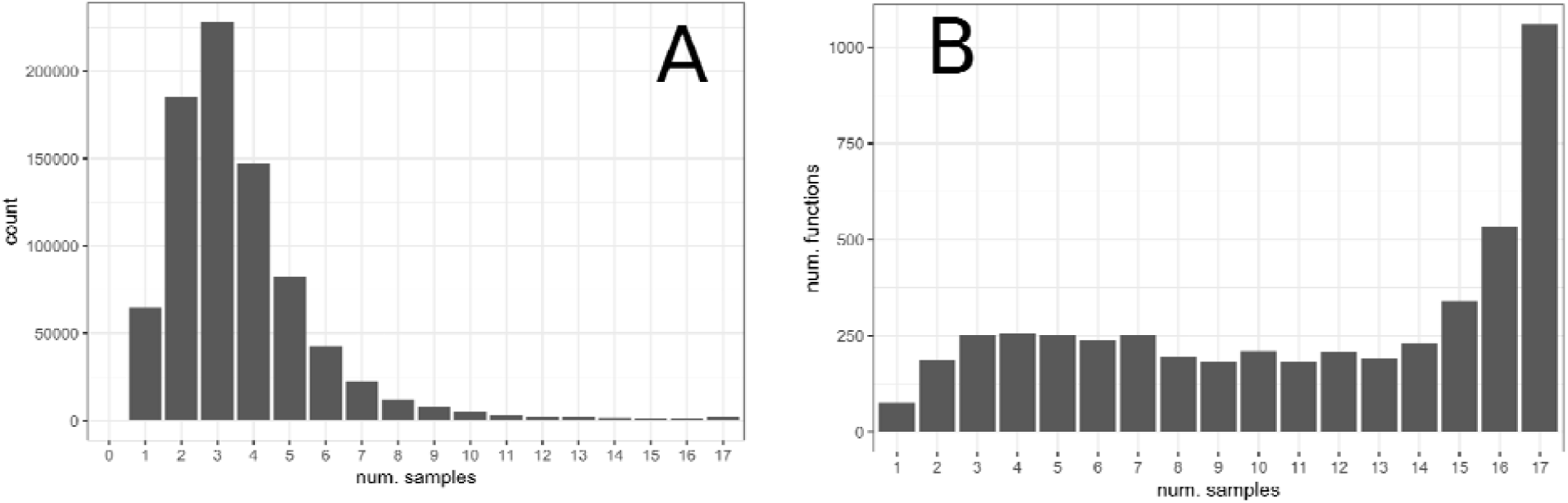
Distribution of transcripts and gene functions across RNA-seq samples. (A) Sample-specificity of assembled transcripts. Each bar denotes the number of transcripts observed in exactly k samples, where k is denoted on the x axis. (B) Sample-specificity of gene functions. Same as in A after collapsing transcripts to unique functional groups (unique IDs from the SEED database).

